# A Pentatricopeptide Repeat Protein in the *Plasmodium* apicoplast is essential and shows sequence-specific RNA binding

**DOI:** 10.1101/388728

**Authors:** Joanna L. Hicks, Imen Lassadi, Emma Carpenter, Madeleine Eno, Alexandros Vardakis, Ross F. Waller, Christopher J. Howe, R. Ellen R. Nisbet

## Abstract

The malaria parasite *Plasmodium* and other apicomplexans such as *Toxoplasma* evolved from photosynthetic organisms and contain an essential, remnant plastid termed the apicoplast. Transcription of the apicoplast genome is polycistronic with extensive RNA processing. Little is known about the mechanism of post-transcriptional processing. In plant chloroplasts, post-transcriptional RNA processing is controlled by multiple pentatricopeptide repeat (PPR) proteins. Here, we present the biochemical characterisation of the single apicoplast-targeted PPR protein. Apicoplast PPR1 is essential, and binds specific RNA sequences corresponding with previously characterized RNA processing sites. We identify the specific binding motif of PPR1. In RNAse protection assays, PPR1 shields apicoplast transcripts from ribonuclease degradation. Our results show that apicoplast RNA processing is under the control of a single protein, thus presenting an Achilles’ heel for the development of new anti-malarial drugs.

## INTRODUCTION

The malaria parasite *Plasmodium falciparum* and related apicomplexan parasites such as *Toxoplasma* evolved from photosynthetic organisms and contain a remnant plastid known as an apicoplast (Gardner, Williamson, & Wilson, 1991; Howe, 1992; McFadden, Reith, Munholland, & Lang-Unnasch, 1996). The ability to photosynthesise has been lost yet the apicoplast remains essential for parasite survival. The apicoplast genome encodes 30 proteins, two rRNAs and 25 tRNAs (Wilson et al., 1996). Primary RNA transcripts are polycistronic and there must be extensive RNA processing to produce individual tRNA, rRNA and mRNA molecules. Many mRNA transcripts show discrete ends coinciding with tRNA sequences, while others show heterogeneous ends within protein coding sequences (R. Ellen R. Nisbet, Kurniawan, Bowers, & Howe, 2016; R. E. R. Nisbet & McKenzie, 2016). The processing of polycistronic transcripts to individual RNAs requires both site recognition and RNA cleavage.

In plants, the primary agents through which the nucleus exerts control on organelle gene expression are pentatricopeptide repeat (PPR) proteins. PPR proteins are encoded in the nuclear genome and are targeted to the mitochondrion or plastid (Barkan & Small, 2014). Plants contain many hundreds of PPRs (Lurin et al., 2004). By contrast, genomes of algae and non-photosynthetic eukaryotes encode relatively few PPR proteins (Manna, 2015; Tourasse, Choquet, & Vallon, 2013). PPR proteins are involved in all aspects of organelle RNA biology, including splicing, editing, transcript stability and translation. In plants, chloroplast PPR mutants show defects in fertility and embryo and seed development (Bryant, Lloyd, Sweeney, Myouga, & Meinke, 2011; Lurin et al., 2004; Prikryl, Rojas, Schuster, & Barkan, 2011; Sosso, Canut, et al., 2012; Sosso, Mbelo, et al., 2012). Additionally, a mitochondrial mutation which causes cytoplasmic male sterility (pollen sterility) is suppressed by a PPR protein (Brown et al., 2003). PPR proteins are sequence-specific RNA-binding proteins. They contain 2-30 tandem repeats, each repeat comprising a 35-amino acid motif (Manna, 2015; Prikryl et al., 2011). The PPR protein family can be divided into two classes: (1) the P-class which contain only the 35 amino acid repeats which stabilize specific RNAs by binding to the 5’ or 3’ termini of RNA transcripts and providing a barrier to exonuclease digestion (Lurin et al., 2004; Pfalz, Bayraktar, Prikryl, & Barkan, 2009; Prikryl et al., 2011; Zhelyazkova et al., 2012); (2) the PLS-class which contains repeats of 31-36 amino acids together with further domains at the carboxyl terminus of the protein. The PLS class can be further divided into E and DYW subclasses, both implicated in RNA editing, carrying out a deamination reaction converting specific cytidines to uridine in plant organelles.

Very little is known about the molecular mechanisms of post-transcriptional processing in the apicoplast. A number of nucleus-encoded, apicoplast-targeted proteins have been identified which may function in RNA processing. Only one RNA-binding protein (*P. vivax PVX_084415*) has been partially characterized, although the stability of the heterologously expressed protein was such that it was not possible to carry out functional assays, though it did bind to uridine rich RNA (García-Mauriño et al., 2018).

Here, we report the identification of a single apicoplast PPR protein. We show that this protein, designated PPR1, is localized within the apicoplast of both *Plasmodium falciparum* and *Toxoplasma gondii*. It is essential for growth of *T. gondii.* Biochemical characterisation of the *P. falciparum* PPR protein shows it binds to a specific RNA sequence and protects RNA transcripts from degradation by ribonucleases. Although the presence of a PPR proteins in the apicoplast is not unexpected, the dependence of a plastid on just a single PPR protein is unique and identifies PPR1 as a good potential drug target.

## RESULTS

### A single apicoplast PPR protein present in both Plasmodium and Toxoplasma

Searches of the *P. falciparum* genome for genes encoding PPR proteins identified only two genes, *Pf*PPR1 (PF3D7_1406400 (PF14_0061)) and *Pf*PPR2 (PF3D7_1233300 (PFL1605W)). Both genes encode proteins with 10 PPR motifs, as predicted by TPRpred (Karpenahalli, Lupas, & Söding, 2007). *Pf*PPR1 belongs to the P-class of PPR proteins, as the repeats all comprise 35 amino acids, and the final PPR motif is situated at the C-terminus of the protein. *Pf*PPR2 may belong to the PLS class, as its PPR elements are not located at its C-terminus. Orthologues of both *Pf*PPR1 and *Pf*PPR2 were found in all *Plasmodium* species with no evidence of paralogues created by lineage-specific gene duplications.

For a protein to be targeted to the apicoplast it must contain both a signal peptide and a plastid-targeting sequence. *Pf*PPR1 and *Pf*PPR2 were analyzed by PlasmoAP and PlasMit for putative apicoplast and/or mitochondrial localization signals (Bender, van Dooren, Ralph, McFadden, & Schneider, 2003; Foth et al., 2003). *Pf*PPR1 analysis by PlasmoAP resulted in 3/4 positive tests for a signal peptide and 5/5 positive tests for an apicoplast targeting peptide, while PlasMit gave a prediction of 99% for not being mitochondrial. This is consistent with an overall strong prediction that *Pf*PPR1 traffics to the apicoplast. PlasmoAP analyses of PPR1 sequences from other *Plasmodium* species similarly predicted that most have an apicoplast localization (Supplementary Table S1), the exceptions being those encoded on genomes with a high GC content, where PlasmoAP is less accurate (Foth et al., 2003). Alignments of PPR1 show that the protein is well conserved across *Plasmodium* species (Figure S1). *Pf*PPR2 was predicted to lack both a signal peptide and a mitochondrial targeting sequence so its location is unknown, but is unlikely to be apicoplast targeted.

To test for the presence of PPRs more broadly in the Apicomplexa we searched for homologues in *Toxoplasma* and *Cryptosporidium.* Analysis of the *Toxoplasma* genome using BLAST and TPRPred, identified five PPR proteins. Only one protein (TGGT1_244050, *Tg*PPR1) contained a predicted signal peptide followed by a plastid-targeting sequence as analyzed by SignalP and iPSORT (Bannai, Tamada, Maruyama, Nakai, & Miyano, 2002; Nielsen, 2017). None of the other four proteins was predicted to include a signal peptide. This indicates that there is only one apicoplast targeted PPR protein in *Toxoplasma*, as is the case in *Plasmodium* spp. The apicomplexan *Cryptosporidium*, which has lost the apicoplast, did not contain any genes encoding PPR proteins.

To test for the localization of the *Pf*PPR1 protein, we expressed recombinant *Pf*PPR1 in *Escherichia coli*, raised poly-clonal antisera in rabbits and used immunofluorescence microscopy to locate this protein. The signal from cells stained with anti-*Pf*PPR1 co-localized with apicoplast-located GFP in the *P. falciparum* D10-ACP_L_ parasite line (Waller, Reed, Cowman, & McFadden, 2000) (Figure 1). This confirmed the predicted apicoplast-localization of *Pf*PPR1. A Western blot of *P. falciparum* lysate probed with the anti-*Pf*PPR1 antibody showed no detectable band, presumably due to low expression of the endogenous protein.

**Figure 1.**
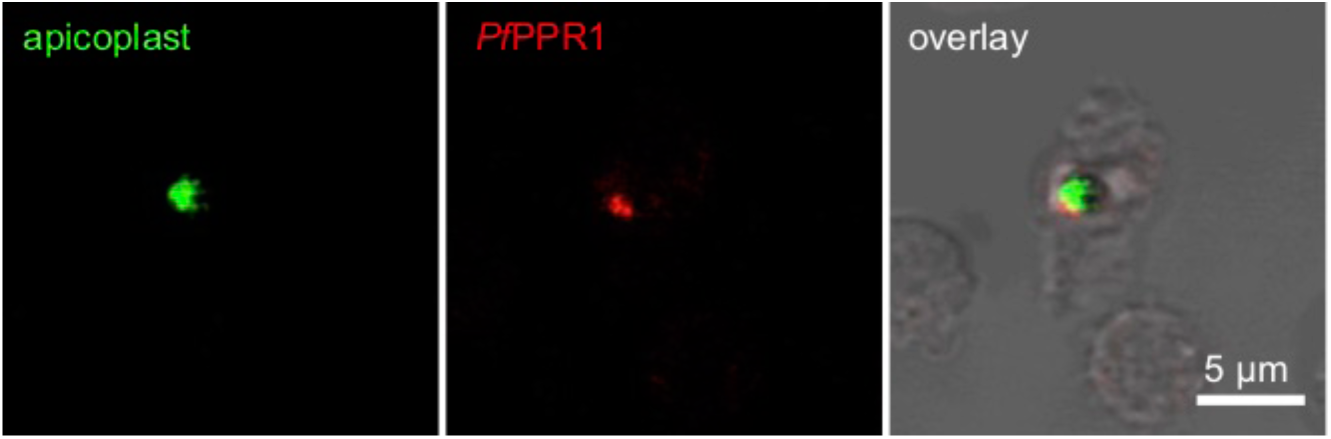
*Pf*PPR1 is localized to the *P. falciparum* 3D7 apicoplast. Immunofluorescence microscopy using *P. falciparum* D10 ACP_L_-GFP parasites that target GFP to the apicoplast (left panel) and an antibody specific for *Pf*PPR1 with a secondary AlexaFluor-568 antibody (middle panel) showed localization of *Pf*PPR1 to the apicoplast (indicated by overlay with bright field image in the right panel).

To test for the localization of the *Toxoplasma Tg*PPR1, a 3′-PPR1-mCherry fusion construct was created to tag the endogenous *Tg*PPR gene. A Western blot probed with antibodies to the mCherry reporter protein showed a faint band, corresponding to low expression levels of the mature fusion protein (Figure 2A, *Tg*PPR1-mCherry), but no signal was apparent by fluorescence microscopy. When the endogenous PPR promoter was replaced by the inducible t7s4 promoter, in cells designated iΔ*Tg*PPR1-mCherry, a higher expression level of this PPR-fusion was seen by Western blot. Furthermore, two bands were present, of apparent sizes consistent with the mature protein and a preprocessed PPR targeting intermediate still bearing the predicted apicoplast targeting peptide (Figure 2A). The presence of these two bands is characteristic for many apicoplast-targeted proteins (Waller et al., 2000). Given the presence of only the shorter, processed band when PPR1-mCherry fusion was expressed from the native promoter, this would indicate near-complete processing under normal expression levels.

**Figure 2.**
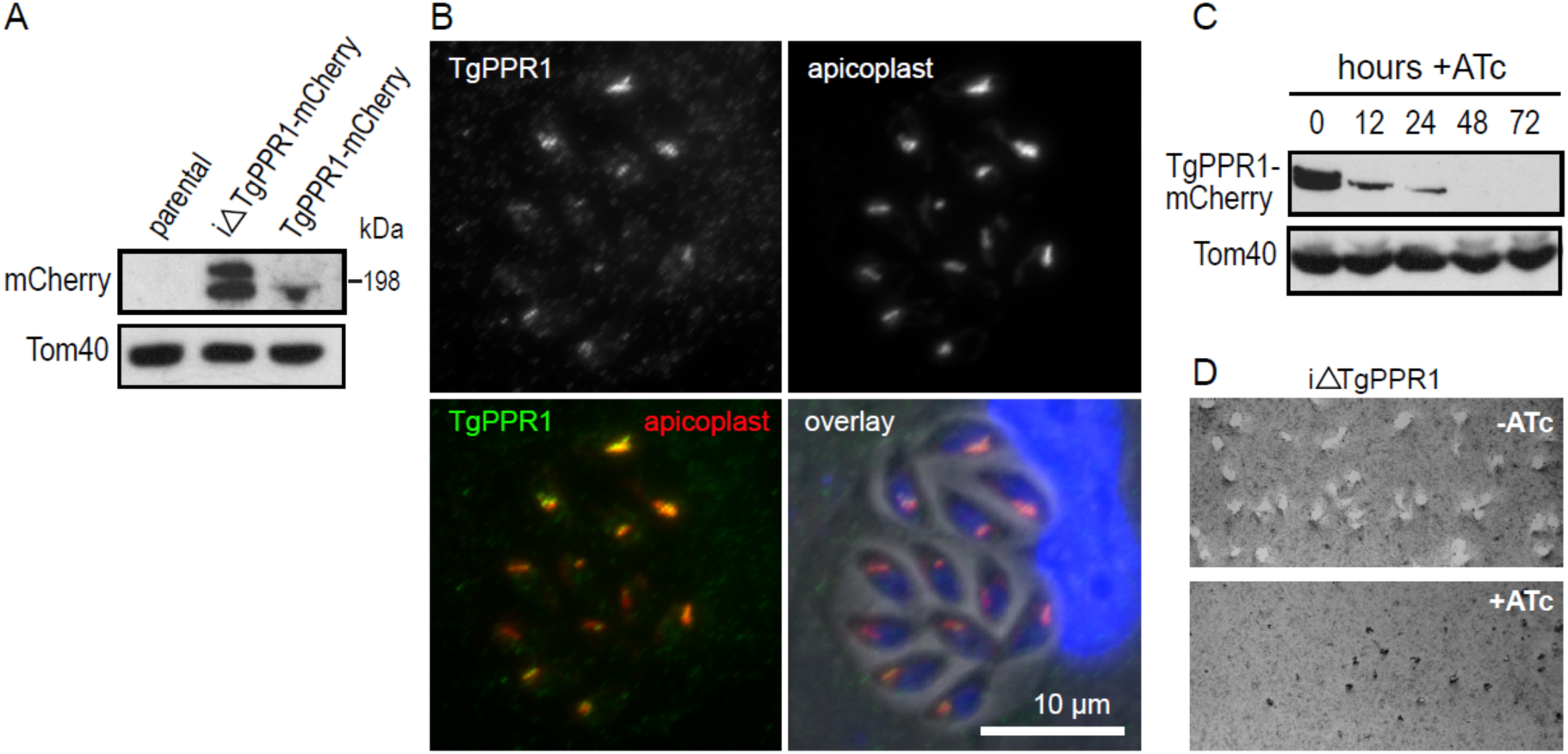
Apicoplast protein *Tg*PPR1 is necessary for parasite growth. (A) Western blot detection of mCherry-tagged endogenous *Tg*PPR1 using either the t7s4 promoter (iΔ*Tg*PPR1-mCherry) or the native promoter (*Tg*PPR1-mCherry). Tom40 acts as a loading control. The presence of two bands in the iΔ*Tg*PPR1-mCherry lane is consistent with a pre-processed PPR targeting intermediate still bearing the predicted apicoplast targeting peptide and mature PPR-mCherry fusion protein. The position of the 198 kDa standard is shown. (B) Co-location of *Tg*PPR1-mCherry expressed from the t7s4 promoter with resident apicoplast biotinylated proteins visualized by streptavidin-staining. DNA staining in blue, *Tg*PPR1-mCherry in green and streptavidin-stained apicoplast in red. (C) ATc-induced knockdown of *Tg*PPR1 assayed over 72 hours. Tom40 acts as a loading control. (D) Eight-day plaque assay shows normal plaque formation in iΔ*Tg*PPR1 cells without ATc-induced *Tg*PPR1 depletion, but no plaques with ATc treatment, indicating that PPR is essential for normal growth.

To confirm apicoplast localization, we performed an immunofluorescence assay on the iΔPPR1-mCherry cells (green) co-stained with streptavidin-594 (red) which serves as an apicoplast marker due to endogenous biotinylated apicoplast proteins (Chen et al., 2015). Using this cell line we could detect *Tg*PPR1-mCherry location and observed it co-locating with the apicoplast steptavidin marker (Figure 2B). We conclude that the PPR protein is localized to the apicoplast, and is normally expressed at a very low level.

### PPR1 is essential for normal growth

As PPR proteins in plants are known to be essential for chloroplast function, we tested if the apicoplast PPR1 was also important for parasite growth. Knock-down of *Tg*PPR1 in the iΔ*Tg*PPR1-mCherry line is induced by the addition of ATc, which represses the t7s4 promoter required for PPR1 expression. ATc treatment of iΔ*Tg*PPR1-mCherry showed rapid depletion of *Tg*PPR1-mCherry with the preprocessed protein undetectable within 12 hours of treatment, and no protein detected after 48 hours by Western blot (Figure 2C). To test for a growth phenotype with PPR1 depletion we used a iΔ*Tg*PPR1 cell line (i.e. t7s4 promoter and no mCherry fusion). Without ATc-induced depletion these cells showed normal growth by an eight-day plaque assay. With ATc treatment no plaques were observed indicating a strong growth inhibition phenotype in cells depleted of *Tg*PPR1 (Figure 2D). This same growth inhibition phenotype was also seen for iΔ*Tg*PPR-mCherry with ATc treatment (not shown).

These results are consistent with *Tg*PPR1 gene disruption being reported to have a negative growth phenotype in a genome-wide CRISPR knockout screen (Sidik et al., 2016). Similarly, in *Plasmodium berghei* a recent genetic screen showed that the PPR1 orthologue (<u>PBANKA_1035800</u>) is also essential to blood-stage growth (Bushell et al., 2017). Together these data suggest that the apicoplast PPR1 is broadly essential to apicomplexan parasites.

### PfPPR1 binds in vitro transcribed apicoplast transcripts

We then tested if *Pf*PPR1 would bind apicoplast RNA transcripts. The recombinant *Pf*PPR1 was assessed for folding by circular dichroism which revealed a folded, alpha helical protein consistent with the alpha helical nature of PPR proteins (Supplementary Figure S2). The protein eluted as a dimer from a gel filtration column (see below), and this dimerization was confirmed by analytical ultracentrifugation following cleavage of the TRX-His_6_ tag by HRV 3C protease (Supplementary Figures S3 and S4). The observed folding and dimerization is consistent with other reported plant PPR proteins (Barkan et al., 2012; Ke et al., 2013) and indicates an appropriate conformation of our purified *Pf*PPR1.

Based on the results of Nisbet et al (R. Ellen R. Nisbet et al., 2016), we generated apicoplast RNA *in vitro* transcripts spanning the *tufA* to *clpC* region of the apicoplast genome as this shows two clearly defined processing sites (at the tRNA-Phe and tRNA-Trp genes), and a second RNA transcript spanning the *LSUrRNA* to *rpoB* region that shows a processing site at tRNA-Thr. To test for *Pf*PPR1-binding we biotinylated the 3’ end of each transcript and performed pull-down experiments against the *Pf*PPR1 protein. *Pf*PPR1 was observed to bind to both transcripts, Figure 3A. As a control, PPR protein was replaced by a DNA-binding protein from *Mycobacterium smegmatis* (AmtR), which had been expressed and isolated using the same procedure as *Pf*PPR1 (Petridis et al.). No binding to AmtR was seen.

**Figure 3.**
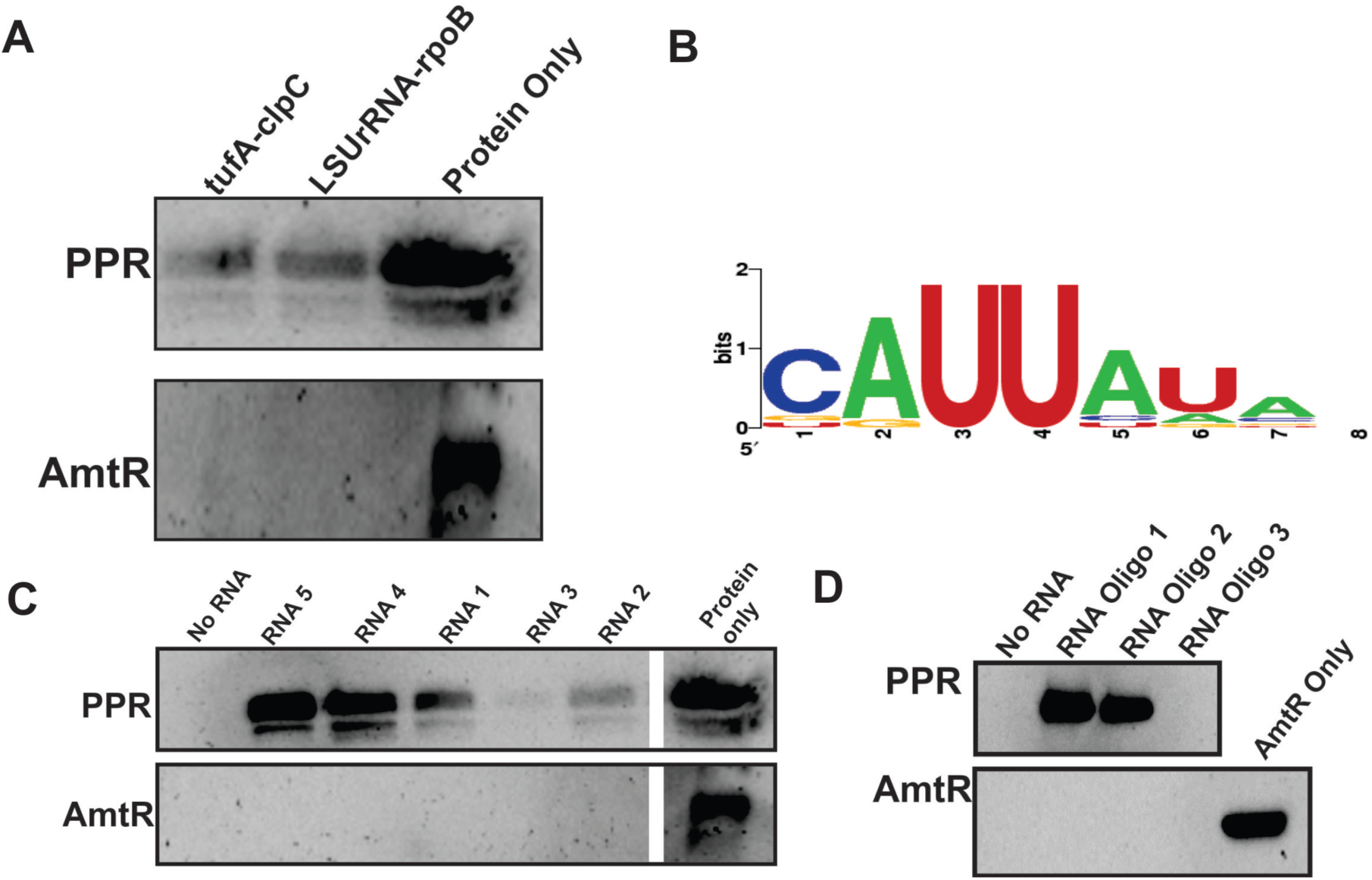
*Pf*PPR1 binds RNA. (**A**) Pull-down assay showing that *in vitro* transcribed apicoplast RNA transcripts (*tufA-clpC* and *LSUrRNA-rpoB*) binds to *Pf*PPR1, as shown by Western blot analysis**. (B)** Weblogo of sequences enriched by SELEX, the height of the letter corresponds to the frequency of that nucleotide at that position. **(C)** RNA transcribed from five clones identified from the final SELEX round were used in a pull-down experiment. Each 150 nt RNA molecule contained a constant 125nt region (not shown) and a variable 25 nt region (shown below). Consensus sequences and sequences with slight variation from the consensus are underlined. As a control, RNA pull-downs were also performed with AmtR. Loading controls are shown to the right of the gel. **(D)** *Pf*PPR1 pull-downs with biotinylated RNA oligonucleotides (sequence shown below) each containing the consensus sequence followed by either a U (RNA Oligo 1, underlined) or an A (RNA Oligo 2, underlined) and a randomly generated six nucleotide sequence (underlined in RNA Oligo 3) using an anti-His antibody demonstrated specificity of PPR for the consensus sequence. The control protein AmtR did not interact with any of the RNA oligonucleotides tested.

**RNA1** <u>UUAA</u>ACUUGAUGCCCGGCGUUUCAG

**RNA2** <u>UUAC</u>CGCGCGUAACACCGGGCCUGU

**RNA3** UUCGGCGACGGAAAGAGUGAAUCCG

**RNA4** UUGUAU<u>UUAU</u>UUAAAAAA<u>UUAU</u>GU

**RNA5** <u>UUAU</u>AACUCGCCUAGACGGGA<u>UUAU</u>

**RNA oligo 1** ACGACA<u>UUAUAU</u>GGUCGGA

**RNA oligo 2** ACGACA<u>UUAUAA</u>GGUCGGA

**RNA oligo 3** ACGACA<u>UGACGA</u>GGUCGGA

### PfPPR1 shows RNA Sequence-Specific Binding

We next sought to determine if *Pf*PPR1 had a sequence-specific preference for RNA-binding. Analysis of the *Pf*PPR1 amino acid sequence using programs designed for the prediction of plant PPR RNA-binding sequences (Barkan et al., 2012; Takenaka, Zehrmann, Brennicke, & Graichen, 2013) did not result in any sequence predictions, presumably due to low sequence identity between plant and apicoplast PPR proteins. To determine any RNA sequence specificity of *Pf*PPR1, we performed SELEX (Systematic Evolution of Ligands by EXponential enrichment) (Manley, 2013). We constructed a SELEX library with a random 25 nucleotide sequence (N25) in the middle of a 150 nucleotide RNA sequence (Manley, 2013). After four rounds of selection using recombinant His_6_-TRX-*Pf*PPR1, sequences containing the motif UUAU were identified in 20 of the 50 final round clones (Figure 3B), with little sequence homology amongst the other 30 final round clones, suggesting a binding motif preferred by *Pf*PPR1. This sequence motif identified by SELEX is very similar to the UUAU apicoplast RNA processing site previously identified (R. Ellen R. Nisbet et al., 2016).

To confirm that *Pf*PPR1 binds the RNA molecules containing the identified sequence motif, we performed PPR pull-down assays using a range of biotinylated 150 nucleotide RNA molecules as ‘bait’ for protein binding. The RNAs were obtained from *in vitro* transcription from five clones isolated in the final round of the SELEX experiment above. RNAs 1, 4 and 5 contained either one or two predicted PPR binding sites while RNA 2 contained a variation of the binding site and RNA 3 lacked the binding site, as shown in Figure 3C. The biotin-labelled transcripts were bound to streptavidin magnetic resin and incubated with recombinant His_6_-TRX-*Pf*PPR1 in a 2:1 protein:RNA molar ratio (as it has been reported to act as a dimer in plants). Western blots were used to detect PPR protein bound to the ‘bait’ RNA. RNAs 4 and 5 showed strongest bound *Pf*PPR1 and each contained two of the consensus binding motifs (Figure 3C). RNAs 1 and 2 that contain slight variations of the consensus sites, UUAA and UUAC respectively, and showed less bound *Pf*PPR1. RNA 3 did not contain the consensus site and exhibited very weak binding to PfPPR1 (Figure 3C). Thus, PfPPR1 binding correlated with presence and number of the consensus binding motif. As a control, the PPR protein was replaced by the *M. smegmatis* DNA binding protein (AmtR (Petridis et al.)) which showed no binding to the RNA transcripts (Figure 3C). We also repeated the experiment with the *Pf*PPR1 protein minus the His_6_-TRX tag and with an MBP tag instead of TRX to ensure that the tag on the PPR protein did not interfere with RNA binding, and no difference in the results was seen (data not shown).

To further test the specificity of *Pf*PPR1 for the UUAU consensus motif, and not elsewhere on the RNA molecules we synthesized three 19 nucleotide RNA oligonucleotides with identical flanking sequences but differing at the potential binding motif. RNA oligos 1 and 2 contained the consensus binding sequence, followed by either an AA or a AU, while oligo 3 did not contain the binding sequence (RNA oligo 1 *<u>UUAU</u>AA*, RNA oligo 2 *<u>UUAU</u>AU*, RNA oligo 3, *UGACGA*). Each RNA oligonucleotide was biotinylated at the 3’ end and used to pull-down *Pf*PPR1 or *M. smegmatis* AmtR (control), as above. Western blot analysis showed that *Pf*PPR1 was recovered from RNA oligos 1 and 2 binding assays, but not using RNA oligo 3. None of the RNA oligos bound to AmtR (Figure 3D). Together, these results demonstrate *Pf*PPR1 shows a strong preference for binding RNA at UUAU.

### PfPPR1 binds RNA as a dimer

PPR proteins THA8 from *Brachypodium distachyon* (Ke et al., 2013), *Arabidopsis thaliana* HCF152 and *Zea mays* PPR10 bind RNA as a dimer (Li et al., 2014; Meierhoff, Felder, Nakamura, Bechtold, & Schuster, 2003; Yin et al., 2013), whereas *Z. mays* PPR4 and PPR5 exist as monomers. We therefore sought to determine if PPR1 bound RNA as a monomer or as a dimer. Analysis by gel filtration chromatography showed the *Pf*PPR1 protein eluted from an analytical gel filtration column at 12.73 ml with a predicted molecular weight of 144 kDa corresponding to a PPR dimer in the absence of any RNA (Figure 4A) and consistent with AUC analysis (Supp Figure 4). To test if this dimer conformation is maintained upon binding to RNA, *Pf*PPR1 was incubated with each of three 150 nt baits used above: RNA 4 and RNA 5 both containing two binding motifs; and RNA 3 that lacks the motif. Incubations were performed in a 1:1 molar ratio at room temperature for 15 minutes. Compared to the no RNA control, RNA 4- and RNA 5-incubated *Pf*PPR1 eluted earlier from the column at 10.97 ml consistent with RNA bound to the *Pf*PPR1 dimer (Figure 4A) (If *Pf*PPR1 bound RNA as a monomer we would expect the elution volume to be greater than 12.73 ml). The presence of the *Pf*PPR1 protein in the elution fraction was confirmed by SDS-PAGE (Figure 4B). The protein fraction was also treated with Proteinase K and RNA extracted by phenol/chloroform treatment. When analyzed via agarose gel electrophoresis, RNA was visible (Figure 4C). These data confirm binding of *Pf*PPR1 to the consensus RNA motif, and that this binding occurs as a PPR dimer.

**Figure 4.**
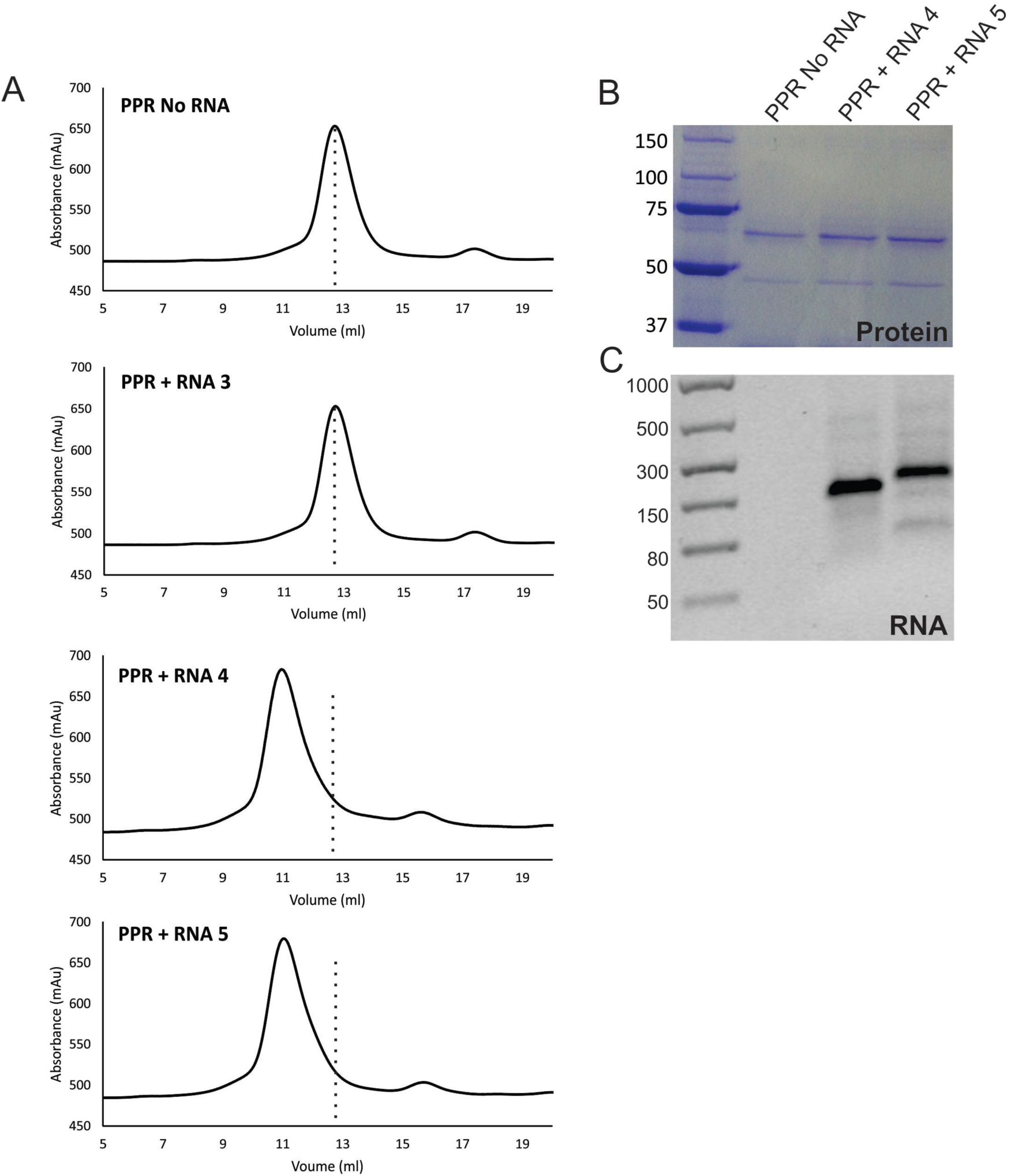
Gel filtration shows a change in elution profile when *Pf*PPR1 is bound to RNA. (A) The elution profiles following gel filtration chromatography of *Pf*PPR1 incubated with RNAs 3, 4 and 5, together with a no RNA control. In each case, a dotted line is shown to indicate where the no RNA control peak is found. (B) Elution peaks for *Pf*PPR1 + no RNA, *Pf*PPR1 + RNA 4 and *Pf*PPR1 + RNA5 were analyzed using SDS-PAGE. The expected molecular weight of *Pf*PPR1-TRX-His_6_ is 72 kDa.(ladder sizes in kDa). (C) Agarose gel electrophoresis of RNA extractions from elution peaks *Pf*PPR1 + no RNA, *Pf*PPR1 + RNA 4 and *Pf*PPR1 + RNA5 (ladder in nt).

### PfPPR1 protects transcripts from ribonuclease activity

As the RNA consensus motif is also associated with known transcript cleavage sites, the binding of *Pf*PPR1 was predicted to be able to protect RNA from degradation by ribonucleases. To test for *Pf*PPR1 protected footprints, we performed RNase protection assays. Three 150 nt RNA molecules with either one or two consensus binding sites (RNAs 1, 4 and 5, as before) were pre-incubated with *Pf*PPR1 protein and then incubated with the RNA endonuclease RNase A. In the absence of *Pf*PPR1 the transcripts were completely degraded by RNase A. However, with pre-incubation with *Pf*PPR1, a small RNA fragment (less than 50 nucleotides) remained after RNase A treatment in each (Figure 5A). We similarly tested for protection of three 19 nucleotide RNA oligonucleotides (RNA oligos 1, 2, and 3, as before). No degradation was evident when the RNA oligonucleotides 1 and 2 were incubated with *Pf*PPR (Figure 5B). In contrast, RNA oligonucleotide 3, which does not contain the binding sequence, was completely degraded by RNase A in the presence of *Pf*PPR1 (Figure 5B). These data show that *Pf*PPR1 protects an RNA region in a consensus binding motif-specific manner.

**Figure 5.**
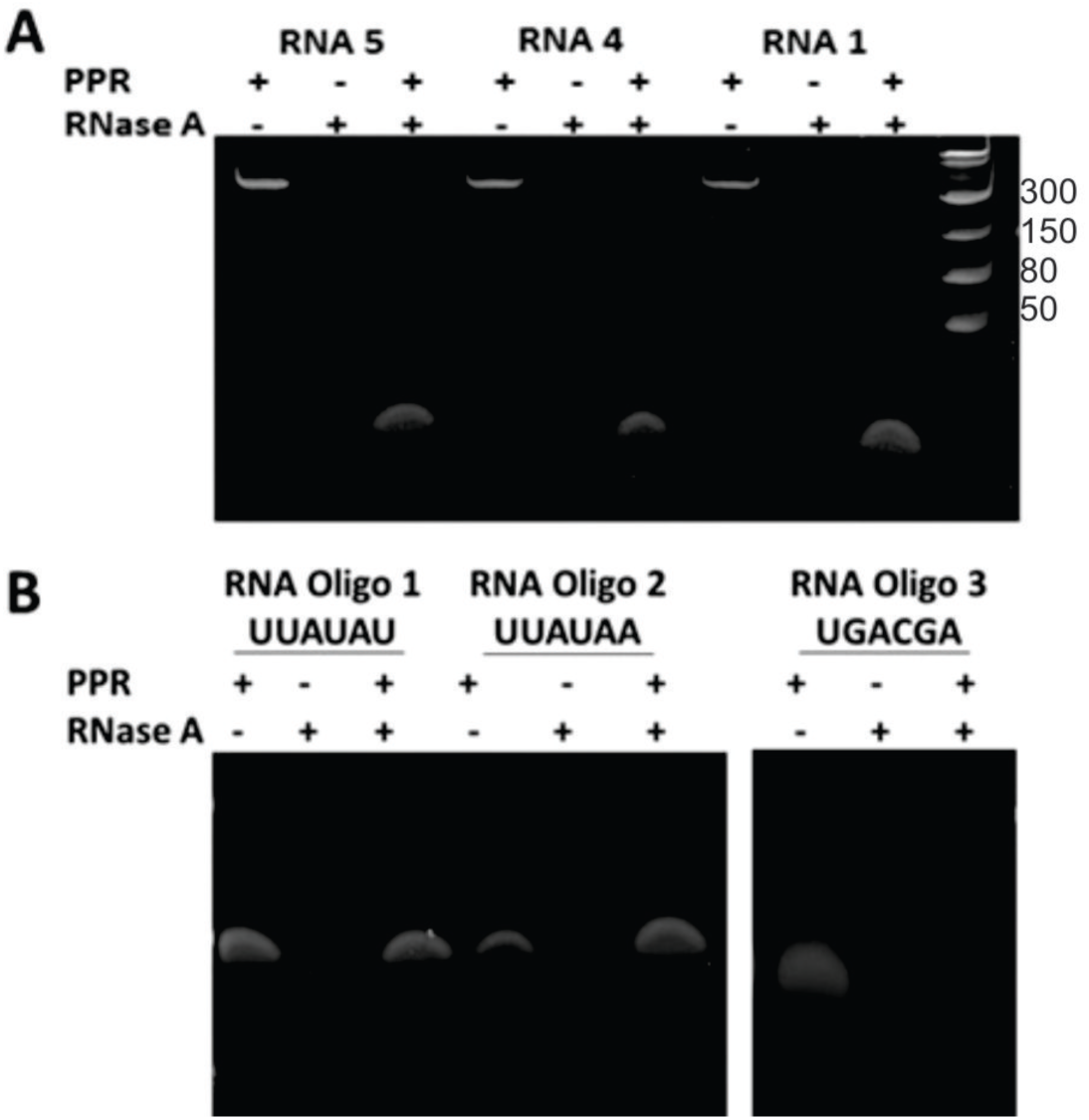
Ribonuclease A protection assays. (A) RNA transcripts 1, 4 and 5 and (B) RNA oligonucleotides 1, 2 and 3 were incubated in a 1:1 molar ratio with *Pf*PPR1 prior to treatment with RNase A. Samples were analyzed using a native acrylamide gel (ladder in nt). Positive controls with no *Pf*PPR1 bound to RNA showed complete degradation by RNase A.

## DISCUSSION

The discovery of a genome of plastid origin in the malaria parasite *Plasmodium* twenty-five years ago was a great surprise. Further investigation revealed the presence of small, but essential organelle subsequently called the apicoplast. We now know a considerable amount about the biochemistry and evolutionary history of this organelle. However, very little is known about how the apicoplast genome itself is transcribed, or how post-transcriptional processing is regulated. Here, we present the characterization of the single apicoplast pentatricopeptide repeat (PPR1) protein and show that it is key to the regulation of post-transcriptional RNA processing.

PPR1 is a nucleus-encoded RNA-binding protein. We confirm that it is targeted to the apicoplast in both *Plasmodium* and *Toxoplasma.* It is essential for normal growth in *Toxoplasma*, and is highly likely also to be essential in *Plasmodium* (Bushell et al., 2017). PPR1 is a P-class PPR protein as it has no additional C-terminal domains and thus no catalytic function. It is predominantly alpha helical in structure, contains 10 PPR motifs, and forms a dimer.

We show that the *Plasmodium* apicoplast PPR1 binds RNA at a UUAU-type motif. This motif is found at known cleavage sites for *Plasmodium* apicoplast transcripts (R. Ellen R. Nisbet et al., 2016). We show that the protein binds *in vitro* to apicoplast RNA transcripts containing known cleavage sites, and can protect RNA from degradation by RNase A. We have previously shown that long, polycistronic RNA molecules are the primary form of transcription in the *Plasmodium* apicoplast (R. Ellen R. Nisbet et al., 2016). These transcripts are processed into individual mRNA, tRNA and rRNA molecules often involving cleavage at the same UUAU motif (R. Ellen R. Nisbet et al., 2016; R. E. R. Nisbet & McKenzie, 2016). Together, these results suggest that the PPR1 protein is involved in the selection of RNA transcript cleavage sites in the apicoplast. The abundance of apicoplast transcripts over the *Plasmodium* erythrocytic life cycle is coordinated with the abundance of *Pf*PPR1 transcripts, with levels of *Pf*PPR1 transcript peaking at the trophozoite stage that immediately precedes the apicoplast transcripts peak at the schizont stage (Bozdech et al., 2003; Le Roch et al., 2003; Llinás, Bozdech, Wong, Adai, & DeRisi, 2006). This indicates that presence of PPR1 is required at the point of increase in apicoplast transcription, and consistent with a role in transcript maturation. Given that *Pf*PPR1 lacks any catalytic domain, and thus is similar to other P-class PPRs, apicoplast PPR1 binding might protect and define mature transcript ends from RNA exonucleases, facilitating their maturation. Alternatively, it could be that *Pf*PPR1 is involved in recruiting the endonuclease activity required to cleave RNA at the specific sites relevant to functional transcript production. In either case, the coordination of PPR1 expression and apicoplast transcription is likely necessary for the spike in apicoplast biosynthetic activity ahead of organelle segregation into schizonts to form the next generation of infective merozoites.

Apicomplexan PPR1 is the first plastid PPR protein to be characterized outside the green chloroplast lineage. Unlike other plant and algal taxa, there is only a single plastid PPR protein in apicomplexan plastids. Apicoplast PPR1, like PPR proteins in plants and green algae, binds at specific RNA sites, known to be the sites of RNA processing. A single and essential apicoplast PPR protein in *Plasmodium* represents a very attractive target for the design of novel antimalarial agents.

## Materials and Methods

### PPR alignment

PPR sequences from *Plasmodium* species were obtained by pBLAST analysis. The PF14_0061 protein alignment was generated in Geneious using a ClustalW algorithm with a BLOSUM cost matrix, a gap open cost of 10 and a gap extend cost of 0.1.

### P. falciparum culture

Blood stage *P. falciparum* D10 ACP-GFP (MRA-568) was cultured according to (Tarr, Nisbet, & Howe, 2011). All work was carried out in accordance with the UK Human Tissue Act 2004.

### Expression of recombinant PfPPR1

PF14_0061 (*Pf*PPR1) was codon optimized for *E. coli* and synthesized by GeneArt and cloned into the pOPIN vector system using InFusion (Takara Biotech) (Berrow et al., 2007). Expression was carried out in BL21(DE3)pLysS in ZY-5052 auto-induction medium (100 μg/ml ampicillin and 34 μg/ml chloramphenicol. PPR1 was putified via HisTrap column on an AKTA FPLC (GE Healthcare). The protein was eluted from the column using an imidazole gradient. Fractions containing His_6_-TRX-PfPPR1 were pooled and concentrated using a Vivaspin concentrator (10 000 MWCO) before gel filtration chromatography using a S200 10/300 analytical size exclusion column. Fractions containing protein were analyzed by SDS-PAGE, and confirmed by MALDI-TOF MS analysis. The His_6_-TRX tag was cleaved from *Pf*PPR1 by incubation of the His_6_-TRX-PfPPR1 fusion protein with 1% (v/v) recombinant HRV 3C protease followed by by gel filtration chromatography. Full details are given in Supplementary methods.

### PfPPR1 antibody production and purification

Recombinant *Pf*PPR1 minus the His_6_-TRX tag was used to generate *Pf*PPR1 antibodies in two rabbits by Pacific Immunology. Pre-immune serum was taken followed by injection of recombinant *Pf*PPR1 plus adjuvant. Four production bleeds were taken at 2 week intervals followed by a final bleed after 3 months.

To purify the anti-*Pf*PPR1 antibody, 150 μl His affinity resin was washed with 1 ml water and three times with 1 x TBS pH 7.6. 250 μl of His_6_-TRX-*Pf*PPR1 (1.7 mg/ml) was added and incubated at room temperature for 15 minutes with agitation. The supernatant was removed and the resin washed three times in 1 x TBS. 1 ml of antibody serum was added and incubated for one hour at room temperature with gentle mixing. The supernatant was removed and the resin washed four times in 1 x TBS. The bound anti-*Pf*PPR1 antibody was eluted by addition of 200 μl 0.1 M glycine pH 2.5. The supernatant was neutralized by the addition of 20 μl 1 M Tris.HCl pH 8.5 to produce purified antibody.

### Immunofluorescence Microscopy for Localization of PfPPR1

Asynchronous *P. falciparum* D10 ACP_L_-GFP cultures were used for immunofluorescence microscopy experiments, essentially following (Tonkin et al., 2004). Purified anti-*Pf*PPR1 antibody was diluted 1:1000 with blocking solution and AlexaFluor-568 Donkey anti-rabbit IgG was diluted 1:2000 with blocking solution. Slides were visualized using an Olympus IX81 confocal microscope at 60 x magnification. Two channels, one to detect GFP fluorescence (eGFP) and the other to detect AlexaFluor-568 fluorescence (Cy3) plus bright field were used to image slides. Images were overlaid using Fluoview version 5.0 microscopy software.

### T. gondii cell culture and generation of cell lines

*T. gondii* RH Δku80/TATi tachyzoites were grown by inoculation in confluent human foreskin fibroblast (HFF) cells as previously described (Striepen, 2007). Endogenous promoter replacement with the t7s4 promoter was induced by Cas9-mediated cleavage at the 5′ end of the *ppr* locus. Plasmid pCRISPR/Cas9-GFP_PPR-sgRNA (see supplemental data) was assembled using the Golden Gate assembly method (Engler et al., 2014). A linear donor molecule including the t7s4 promoter and DHFR resistance gene was amplified from plasmid pPR2-HA3 (Katris et al., 2014) with primers KDPPR-Fwd and KDPPR-Rev that included flanking sequences directed to the 5′ end of the *ppr* locus on either side of the Cas9 cleavage site. pCRISPR/Cas9-GFP_PPR-sgRNA and this linear donor were co-transfected into *T. gondii* TATiDku80 parasites (a kind gift from Lilach Sheiner and Boris Striepen, U. Georgia; (Sheiner et al., 2011)) and transformants selected on pyrimethamine and cloned by limiting dilution(Katris et al., 2014; Striepen, 2007). Successful promoter replacement was verified by PCR from genomic DNA. Endogenous in-frame 5′ tagging of the *ppr* locus with reporter protein gene *mCherry* was achieved using plasmid pPPR-mCherry_CAT (see supplemental data) assembled using the Golden Gate method. Prior to transfection of parasites this plasmid was linearized with *Bam*HI, and transformants were selected with chloramphenicol and cloned by limiting dilutions (Striepen, 2007).

### Toxoplasma PPR assays

Western blot detection of SDS-PAGE resolved whole cell lysates was performed using a rabbit anti-mCherry (1/1000 dilution) (Abcam) and anti-TOM40 as a control (Katris et al., 2014). Immunofluorescence microscopy was performed on intracellular tachyzoites using anti-mCherry (1/1000) with secondary antibody AlexaFluor-488 Goat anti-rabbit IgG (Life technology). Apicoplasts were co-stained with AlexaFluor-594 anti-Steptavidine (Life technology)(Chen et al., 2015). Samples were mounted with ProLong Diamond antifade mountant with DAPI (Invitrogen) and sealed with nail polish. Cells were imaged using an Inverted Nikon Eclipse Ti microscope, a Nikon objective lens (Plan APO, 100x/1.45 oil), and a Hamamatsu C11440, ORCA Flash 4.0 camera.

PPR knockdown was induced by addition of anhydrotetracycline (ATc) (0.5 μg/ml) to the growth medium upon parasite inoculation of HFF cells. For growth assays extracellular parasites were filtered, counted by haemocytometer, and 500 parasites added to 25 cm^2^ tissue culture flasks containing a confluent monolayer of HFF cells. To visualize plaque sizes in the presence or absence of ATc, flasks were aspirated, fixed with 5 ml 100% ethanol (5 minutes), stained with 5 ml crystal violet solution (15 minutes) then washed once with phosphate-buffered saline (PBS) and dried before imaging.

### SELEX for determination of PfPPR1 RNA sequence specificity

A SELEX library was constructed as in (Manley, 2013). SELEX using recombinant His_6_-TRX-*Pf*PPR1 was carried out as per the protocol in (Manley, 2013). Four rounds of selection were performed, following which PCR products were cloned into pGEM-T easy (Promega) and transformed into chemically competent *E. coli* DH5α and plated onto LB agar (100 μg/ml ampicillin, 0.1 mM IPTG and 40 μg/ml X-gal). Plasmids were extracted from 50 clones and sequenced. In addition, 10 input clones were sequenced from each round to determine enrichment.

### In vitro RNA Transcription and 3’ End Biotinylation

Apicoplast *P. falciparum* apicoplast PCR products were obtained using the following primers: LSUrRNA Fwd/rpoB Rev and tufA Fwd. clpC Rev. T7 promoter sequences (TAATACGACTCACTATAG) were added in a further round of PCR with the T7 promoter sequence appended to the 5’ end of the forward primer. Additionally, PCR products were obtained from SELEX clones (above) The Ambion T7 MEGAScript Kit was used for *in vitro* transcription. For 3’ end biotinylation 50 pmol RNA transcript was heated at 85 °C for 3-5 minutes. Once on ice, 3 μl 10 x T4 RNA ligase buffer (NEB), 1 μl rRNasin (Promega), 50 pmol RNA, 1 μl pCp Biotin, 2 μl T4 RNA Ligase (NEB), water to 15 μl and 15 μl 30% PEG was added. Ligation reactions were incubated overnight at 16 °C. 70 μl water was added followed by 100 μl chloroform:isoamyl alcohol (49:1). Reactions were centrifuged at 13 000 x g for three minutes and the upper phase (aqueous layer) removed and transferred to a new tube. 10 μl 3 M sodium acetate pH 5.2 and 250 μl 100% ethanol were added to precipitate RNA and stored at  −20°C. The RNA was pelleted by centrifugation at 13 000 x g, 4 °C for 20 minutes, washed with 70% ethanol and centrifuged again. The pellet was resuspended in 20 μl water and RNA was quantified with a Nanodrop-1000.

### Biotinylated RNA-PfPPR1 Pull-downs

25 μl of resuspended streptavidin magnetic beads (Thermo Scientific) were washed in 1 ml water and then three times in binding buffer (10 mM HEPES pH 7.5, 20 mM KCl, 1 mM MgCl_2_, 1 mM DTT) and resuspended in 20 μl binding buffer. 400 nM biotinylated RNA transcript was added and incubated at 4 °C for 15 minutes. The beads were washed three times in 1 ml binding buffer and resuspended in 20 μl binding buffer. 800 nM His_6_-TRX-*Pf*PPR1 was added followed by incubation at room temperature for 15 minutes with gentle agitation. The beads were washed three times with 1 ml binding buffer and resuspended in 25 μl 4 x SDS loading dye, heated to 100 °C for 5 minutes and loaded onto a 4-15% SDS-PAGE gel (BioRad). Following PAGE, the samples were transferred via wetblot to PVDF membrane for western blot analysis with a mouse-anti His antibody and a secondary goat anti-mouse antibody conjugated with HRP. Western blots were visualized using Western Bright Quantum (Advasnsta) chemiluminiscent substrate and exposed using a CCD camera (Genebox).

### Gel Filtration Chromatography

His_6_-TRX-*Pf*PPR1 was incubated with RNA transcripts in a 1:1 molar ratio in 1 x binding buffer (10 mM HEPES pH 7.5, 20 mM KCl, 1 mM MgCl_2_ and 1 mM DTT) for 15 minutes at room temperature. A control without RNA was included in these reactions. Samples were analyzed using a S200 10/300 analytical gel filtration column pre-equilibrated in 50 mM Tris pH 8.0, 200 mM NaCl, 5 mM DTT.

### RNase A protection assays

10 μM RNA, 1 x binding buffer buffer (10 mM HEPES pH 7.5, 20 mM KCl, 1 mM MgCl_2_ and 1 mM DTT) and 10 μM PfPPR1 in a volume of 20 μl were incubated for 15 minutes at room temperature, followed by addition of 0.01% (v/v) 20 mg/ml RNase A and incubated at 37 °C with shaking for 30 minutes. Reactions were stopped by the addition of 10 μl 2 x formamide loading dye (95% formamide, 0.025% w/v bromophenol blue, 0.025% xylene cyanol FF, 5 mM EDTA) and heating to 70 °C for 5 minutes. Assay reactions were analyzed by 12% urea-denaturing PAGE gel, and visualized with SYBR Safe nucleic acid stain. Three controls (no *Pf*PPR1, no RNA, no RNase) were carried out, where the reagent was replaced by water.

## ADDITIONAL INFORMATION

We thank the numerous anonymous blood donors. The research project was approved by the Human Biology Research Ethics Committee, School of Biological Sciences, University of Cambridge, project 2007.04. All data are included in this article and its supplementary information files. We have no competing interests. This work was funded by a Wellcome Trust project grant (094249) to CJH and RERN, a MRC grant (MR/M011690/1) to RFW, an MRC Confidence in Concept grant to RERN and CJH and a University of Cambridge Returning Carers Award to RERN. All authors gave final approval for publication.

## Author contributions

JLH, IL, EC, ME, AV and RERN performed the experiments. RFW provided expertise in *Toxoplasma.* JLH, RFW, CJH and RERN drafted the manuscript. All authors read and approved the final manuscript.

## ACKNOWLEDGEMENTS

We thank Dr Katherine Stott at the Biophysics facility in the Department of Biochemistry at the University of Cambridge for assistance with CD and AUC experiments, and Dr. Joanna-Marie Howes for assistance with confocal microscopy. We thank Prof Mark Carrington, Dr Erin Butterfield and Dr Richard Dorrell for helpful discussion.

